# Alterations in RNA editing in skeletal muscle following exercise training in individuals with Parkinson’s disease

**DOI:** 10.1101/2023.05.30.542987

**Authors:** Heather Milliken Mercer, Aiswarya Mukundan Nair, Angela Ridgel, Helen Piontkivska

## Abstract

Parkinson’s Disease (PD) is the second most common neurodegenerative disease behind Alzheimer’s Disease, currently affecting more than 10 million people worldwide and 1.5 times more males than females. The progression of PD results in the loss of function due to neurodegeneration and neuroinflammation. The etiology of PD is multifactorial, including both genetic and environmental origins. Here we explored changes in RNA editing, specifically editing through the actions of the Adenosine Deaminases Acting on RNA (ADARs), in the progression of PD. Analysis of ADAR editing of skeletal muscle transcriptomes from PD patients and controls, including those that engaged in a rehabilitative exercise training program revealed significant differences in ADAR editing patterns based on age, disease status, and following rehabilitative exercise. Further, deleterious editing events in protein coding regions were identified in multiple genes with known associations to PD pathogenesis. Our findings of differential ADAR editing complement findings of changes in transcriptional network identified by a recent Lavin et al. (2020) study and offer insights into dynamic ADAR editing changes associated with PD pathogenesis.

## Introduction

Parkinson’s Disease (PD) is the most common motor neurodegenerative disease with disability and death due to PD increasing faster than any other neurological disorder (1). The characteristic symptoms of PD including tremor, troubles with movement and walking, and a mask-like facial expression are largely due to the loss of dopaminergic neurons in the substantia nigra pars compacta. While the effects of PD within the central nervous system (CNS) are well documented (2–5), the resulting impacts on skeletal muscle are not fully understood (6). Several studies have shown that high-intensity resistance training increases muscle strength and improves symptoms in individuals with PD (7–12). A recent study (6) showed that high-intensity resistance training resulted in differential gene expression in skeletal muscle samples in individuals with PD after completion of a 16 week intervention. These changes in transcriptional networks were correlated with increases in muscle mass and strength, improved cognition, feelings of well-being, scores on the Unified Parkinson’s Disease Rating Score (UPDRS) and the Parkinson’s Disease Questionnaire (PDQ-39) (6). Further, the decreased neuronal signaling observed in the brains of PD patients was positively improved following exercise as evidenced by the increase in the amplitude of low-frequency fluctuations (ALFFs) within the right ventromedial prefrontal cortex (PFC), left ventrolateral PFC, and bilaterally within the substantia nigra (SN)(13). These data show that exercise not only affects the cellular biology within the skeletal muscle of PD patients, but also initiates changes in the CNS.

While many studies have focused on differential gene expression in the progression of PD (14–18), other genetic influences may factor into the neuromuscular manifestation and pathogenesis of PD. RNA editing by Adenosine Deaminases Acting on RNA (ADARs), which causes adenosine to inosine deamination, is the most common mechanism of post-transcriptional RNA editing in the nervous system and integral to neurodevelopment (19–22). Unlike gene mutations, which result in permanent changes to the genome, RNA editing can be dynamically and differentially regulated and controlled spatiotemporally in a nuanced manner (23,24). Patterns of ADAR editing vary among species in terms of editing levels, targets, and ADAR isoforms, with the vast majority of editing occurring in non-coding regions of the genome (25–27). Two mammalian loci, ADAR1 (ADAR) and ADAR2 (ADARB1), are catalytically active and expressed in many tissues. In addition, ADAR1 has an interferon-inducible isoform, underlying a connection between the immune system activation and RNA editing (28). ADAR3 (ADARB2) does not show catalytic activity and is only found in the brain, however it is thought to be important during neurodevelopment as an editing inhibitor (22). While post-transcriptional RNA editing is widespread, its role in skeletal muscle is not completely understood. Hsieh et al. (29) suggests that RNA editing is integral to the process of myogenesis although other studies have inferred that the editing level is relatively low in skeletal muscle compared to other types of tissue(30,31). These seemingly contradictory findings suggest potential dynamic relationships between RNA editing in myogenesis and global editing rates.

Although the majority of ADAR editing occurs in non-coding regions (25), ADAR editing also affects protein coding sequences. Albeit lower in numbers, RNA editing sites in protein coding regions and miRNAs are relatively conserved in mammals and can lead to recoding events (27,32–36). Notably, recoding events can have profound phenotypic consequences due to a mere single nucleotide change within a key amino acid encoding codon. ADAR editing has been implicated in the development and progression of multiple neurological, neurodegenerative and psychiatric disorders such as epilepsy (37–40), autism (41,42), schizophrenia (43), Amyotrophic Lateral Sclerosis (ALS) (44–47), Huntington’s Disease (HD) (43), Alzheimer’s Disease (AD) (43,48–50), schizophrenia (51–56), suicide (51,53,54,56), and depression (57,58). Similar to other neurodegenerative diseases, PD is a complicated disease with multiple phenotypes that vary across individuals, as well as by age and sex (59,60). Therefore, it is likely that multiple complicating factors such as the aging process, hormones, genetic and epigenetic factors, including dynamic ADAR editing, may play a role in PD pathology and/or response to exercise.

Utilizing a list of 737 genes (61) (S1A) known to function in PD pathology and a list of 704 genes shown to be differentially expressed in Pre-Training PD and Post-Training PD samples (6) (S1B), we examine whether ADAR editing patterns change due to age and disease state within proteins whose function has been linked with the manifestation and progression of PD. We further compared editing patterns between pre- and post-training PD skeletal muscle samples, to determine whether rehabilitative exercise facilitates changes in ADAR editing. Our results show shifts in ADAR editing patterns between groups and offer insights into dynamic gene regulatory mechanisms underlying PD pathogenesis and response to exercise.

## Materials and Methods

### Transcriptomics dataset

The transcriptomics dataset used in this study included samples from various subgroups from the skeletal muscle RNA-seq study of Lavin et al. (6), namely, samples representing Older Males category (n=9), PD Males (n=9), Younger Males (n=9), and Post-training PD Males (n=4), respectively. Category designations followed Lavin at al. (6) definitions, with Older Males ranged from 50-79 years old, PD Males from 54-76 years old, Younger Males from 23-35 years old, and Post-Training PD Males from 54-76 years old. Pre-training PD sample data (n=4) was included in the PD Male category, when making comparisons to Older Males and Post-Training PD Males but were also used independently for within-patient comparisons to Post-Training PD samples. Post-Training participants completed a 16-week resistance rehabilitative training program, as described in Lavin et al. (6). Briefly, this program consisted of thrice weekly sessions of strength, power, endurance balance, and functional training lasting between 35-45 minutes. Because the Lavin et al. (6) study included only a few female samples (Older Females =3, PD Females =3, Younger Females =3, and Post-Training Females =1), and our prior studies have suggested sex-specific differences in ADAR editing patterns (62), we focused our analyses on male samples. Table 1 lists the pairwise comparisons between analyzed groups.

**Table 1.**
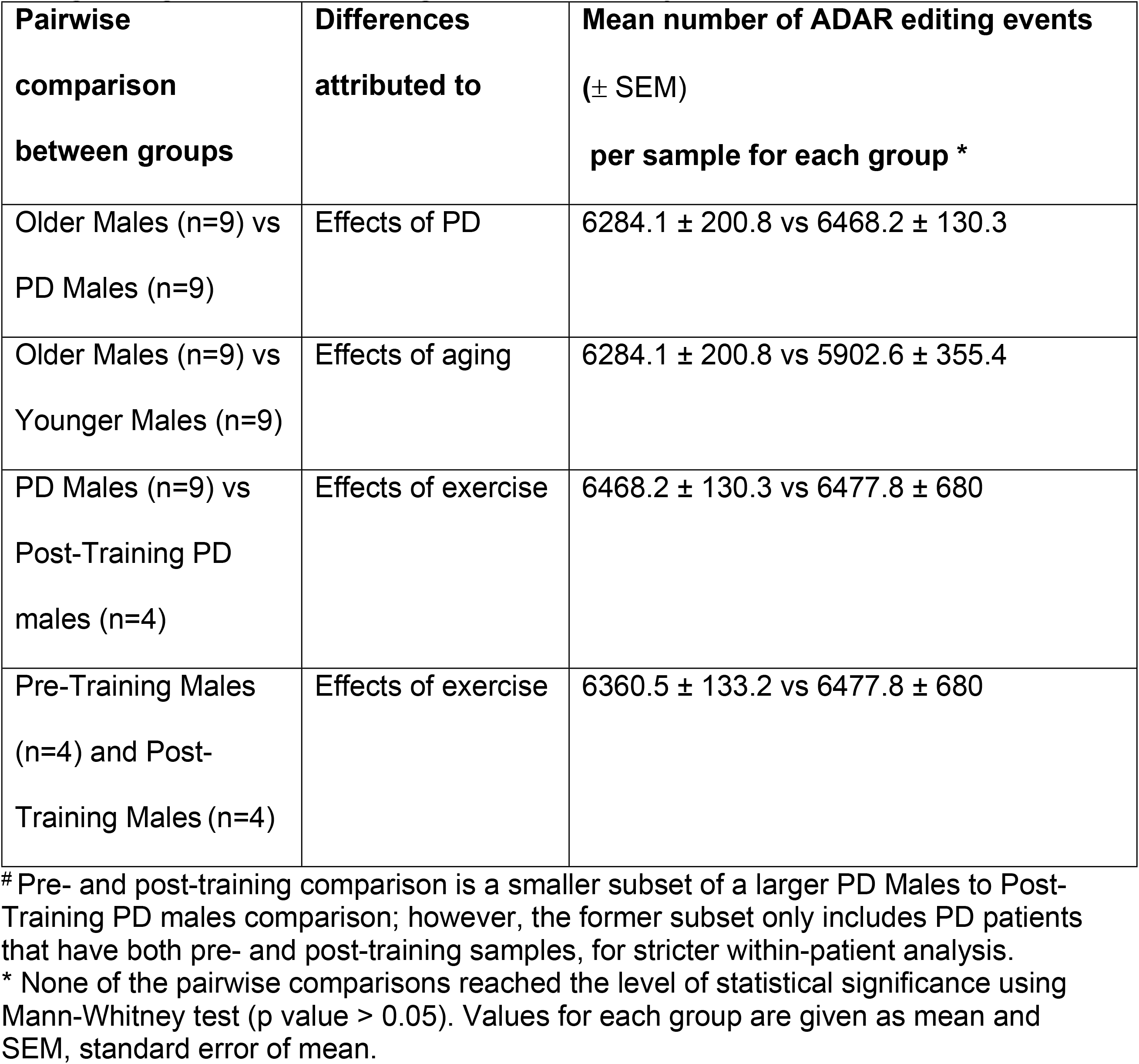
Pairwise comparisons between control and treatment groups, and changes in global ADAR editing parameters if any.

### ADAR editing inferences from RNA-seq data

RNA-seq data from Lavin et al. (6) study was downloaded from NCBI SRA (Bioproject PRJNA588234; GSE140089) (63). Samples were sequenced using paired 50-bp-long reads, with the number of mapped reads varying between ∼33.7 million (33,726,174) and ∼82 million (82,025,036), with an average of ∼59.9 million (59,956,071) reads per sample. Information on mapped reads of samples utilized in this study are shown in S2.

The Automated Isoform Diversity Detector (AIDD) pipeline (64) was used to infer ADAR editing events. Briefly, fastq files of individual patients were trimmed and aligned to the chosen human reference (GRCh37) using HISAT2 (65). Once aligned, the transcriptome was assembled using Stringtie (66), to estimate levels of individual gene expression as Transcripts Per Kilobase Million (TPM). ADAR editing events were inferred using GATK haplotype caller (67) following the best practices, as described in Plonski et al. (64). VCF files are available at https://doi.org/10.5281/zenodo.7971793. SnpEff (68) was used to predicts the effects and functional impacts of predicted edited variants, as described below. Variants identified are putative ADAR editing events described as such based on known ADAR editing sites, but for simplicity will be further described throughout this study as simply “editing events”.

### Inferences of editing consequences

A list of 758 genes with suspected associations to PD were downloaded from NCBI (downloaded April 20, 2022) (61) and converted to Ensembl gene and transcript IDs, resulting in 737 gene and 7827 transcript IDs (S1A) utilizing Ensembl Biomart (69). The NCBI PD Gene List (n=737) is one of the multiple GeneRIF (Gene Relevance into Function) lists, which are created through three primary methods: extraction from published literature by the National Library of Medicine staff, summary reports from HuGE Navigator (70), and user submissions from a Gene record (71).

SnpEff annotations were collated by sample and patient group. NCBI PD genes found within PD, control, and Post-training PD samples, were compared between groups and analyzed for gene overrepresentation within Reactome pathways (72) using the Database for Annotation, Visualization, and Integrated Discovery (DAVID) (73–75)interface. We focused on Reactome pathways because a larger number of our genes of interest (75.8%) had assigned Reactome annotations, compared to KEGG pathways annotations (67.5%). Only genes affected by high or moderate impact editing events occurring in at least 2 samples were considered. High impact editing events include those A-to-G or T-to-C nucleotide changes that result in large duplications or deletions, deleted or duplicated exons, frameshifts, gene deletions, etc. Moderate impact editing events include those that result in exon deletion or duplication, codon deletion or insertion, nonsynonymous coding, etc. High and moderate impact editing events by biotype (protein coding, nonsense mediated decay, etc.) were also considered in this analysis. Significance of overrepresentation is calculated as a p-value with Binomial Test and False Discovery Rate (FDR) using the Benjamini-Hochberg approach (72).

We also used SNPNexus (76) interface for SIFT (Sorting Intolerant from Tolerant) (77) analyses of editing effects on protein function and nonsynonymous coding. Only genes and transcripts with known associations to PD (61), an absence of dbSNP association, and A/G and/or T/C variants were used in this analysis. Editing events occurring in less than 2 samples were removed. Similar to SnpEff annotations, the genes in which deleterious and tolerated editing events occurred were further scrutinized to identify Reactome pathways in which those genes were over-enriched.

Chi-square test was used to compare the numbers of editing events across various categories, such as editing events with high and moderate impact, protein coding, nonsense-mediated decay, deleterious, and tolerated, using GraphPad Prism (version 9.5.1) (78) when categorical data was being analyzed. For ADAR expression, non-parametric Krushkal-Wallis tests were used. For comparing the total number of putative ADAR editing events per sample, as a proxy of global editing levels, non-parametric Mann-Whitney test was used.

## Results

We would first like to note that the implications of our findings should be thought of in terms of dynamic downstream changes to gene regulation within functional (such as Reactome) pathways stemming from identified (putative) ADAR editing events within gene members of these pathways, rather than attempting to categorize such changes into groups where “edited” is synonymous with “disease causing” and “unedited” signifies a “non-disease causing” event. Importantly, both directions of editing events should be considered as editing change; in other words, status of an editing change can be assigned whether the transcript that is normally edited but is no longer edited in a specific condition, or if the transcript becomes edited in a specific circumstance. In fact, many genes experience ADAR editing as part of their normal regulation, and thus, aging is expected to correlate with changes in editing patterns (22,79). Likewise, some genes stop being edited as part of the normal developmental progression (20) or due to physiological changes (24,80,81). Thus, the major challenge is to understand whether or how shifts in editing patterns across multiple genes within a pathway reflect downstream changes in gene regulation and subsequent physiological changes, including the disease process. Thus, in this study we primarily focus on interpreting observed shifts in ADAR editing events in the context of functional consequences to affected genes, including likely changes in gene regulation patterns among impacted pathways.

### Physiological pathways affected by PD genes in which high or moderate impact editing events and deleterious protein outcomes are found vary between PD, Control, Pre- and Post-Training PD samples

SnpEff (68) annotations of putative ADAR editing events were filtered for genes in which high or moderate impact editing events occurred within genes from the NCBI PD Gene List. Entries with high or moderate impact editing events in genes that were found in less than two samples were removed to identify 168 unique genes in Older Male samples, 184 unique genes in PD Male samples, 176 unique genes in Younger Male samples, 113 unique genes in Pre-Training PD samples, and 88 unique genes in Post-Training PD Male samples (S3).

Analysis of Reactome pathways in which these genes were enriched showed that while many of the same pathways were overrepresented among ADAR edited genes across different subject categories, some pathways were uniquely present only in one specific category. We used the DAVID interface to identify Reactome pathways in which PD genes were over-represented including only pathways with at least 10 genes, and at least 10% of genes in the sample gene list associated with the pathway, as well as reported FDR p-value less than 0.05. For consistency, we also included pathways where some patient categories did not meet the “10% of a pathway” criteria if that was met in at least one other patient category (Fig 1).

**Fig 1:**
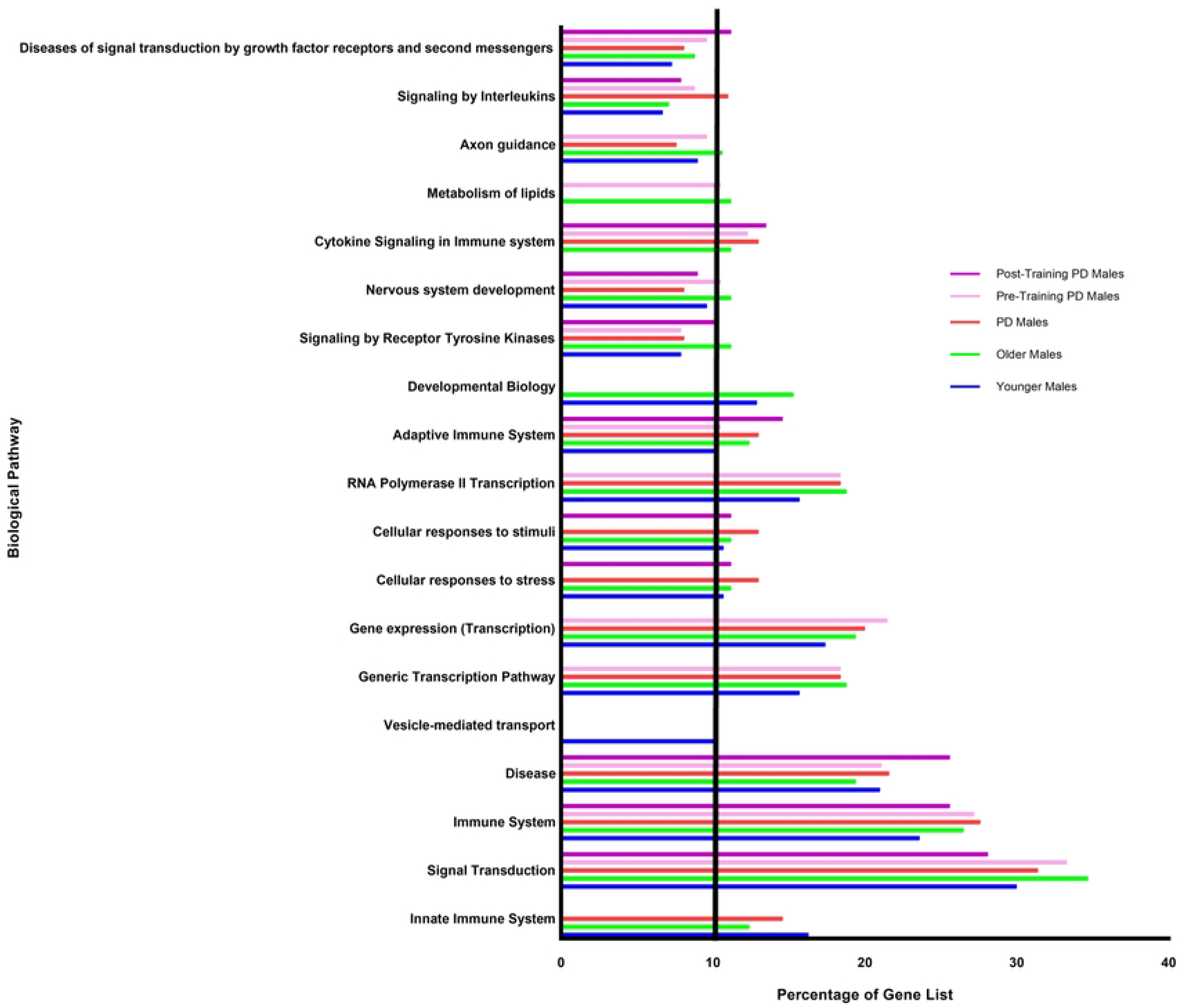
Distribution of ADAR edited genes by overrepresented pathways in each patient category. PD genes in which high or moderate impact editing events occurred within at least 2 samples within a sample group were analyzed via DAVID to identify overrepresented pathways. Pathways where at least one patient category had at least 10% of genes identified as ADAR edited targets are included. Shifts in editing patterns across pathways can be observed between ages, disease status, and training state.

Specifically, edited genes were overrepresented across all sample groups in Signal Transduction, Immune System, Disease, Generic Transcription Pathway, Gene Expression, Adaptive Immune System, Signaling by Receptor Tyrosine Kinase, Nervous System Development, Signaling by Interleukins, and Diseases of Signal Transduction by Growth Factor Receptors and Second Messengers.

When high or moderate impact editing events affecting PD genes are compared between the sample groups, interesting patterns of changes in RNA editing emerge. The proportion of genes showing evidence of high or moderate impact ADAR editing was compared between sample groups to analyze functional relevance (Fig 1). When comparing the ratio of genes devoted to specific functional pathways between Older Males and Younger Males, marked differences were observed in Vesicle Mediated Transport (Older Males 0%, Younger Males 10.1%), Metabolism of Lipids (Older Males 11.2%, Younger Males 0%), and Cytokine Signaling in Immune System (Older Males 11.2%, Younger Males 0%). In Older Males, Developmental Biology was overrepresented while in PD Males, it was not (Older Males 15.3%, PD Males 0%). Metabolism of Lipids pathway also differed between groups (Older Males 11.2%, PD Males 0%). When the pathways in which genes with high or moderate impact edits are over-represented are compared between PD Males and Younger Males, Vesicle Mediated Transport (PD Males 0%, Younger Males 10.1%), Developmental Biology (PD Males 0%, Younger Males 12.9%), and Cytokine Signaling in Immune System (PD Males 13%, Younger Males 0%) varied between the groups. Between Pre-Training PD Males and Post-Training PD Males, differences in pathway over-representation were found between Generic Transcription Pathway (Pre-Training 18.4%, Post-Training 0%), Gene Expression (Pre-Training 21.5%, Post-Training 0%), Cellular Response to Stimuli (Pre-Training 0%, Post-Training 11.2%), Cellular Response to Stress (Pre-Training 0%, Post-Training 11.2%), RNA Polymerase II Transcription (Pre-Training 18.4%, Post-Training 0%), Metabolism of Lipids (Pre-Training 10.5%, Post-Training 0%), and Axon Guidance (Pre-Training 9.6%, Post-Training 0%), but did not differ greatly in the following pathways when the entire PD group was compared to Post-Training samples: Cellular Response to Stimuli (PD Males 13%, Post-Training PD Males 11.2%), Cellular Response to Stress (PD Males 13%, Post-Training PD Males 11.2%), and Metabolism of Lipids (PD Males 0%, Post-Training PD Males 0%). Additionally, when all PD samples are compared to Post-Training PD Males, Innate Immune System pathway is shown to be over-enriched in PD Males, but not in Post-Training PD Males (PD Males 14.6%, Post-Training PD Males 0%).

The Lavin et al. (6) differentially expressed gene list (n=704 unique genes) from skeletal muscle samples was compared to the NCBI PD gene list (61) (n=737), identifying 38 unique genes in common between the two lists, which will be referred to as the Genes of Interest in PD gene list for clarity (S Fig 1). The 38 Genes of Interest in PD genes were compared to 16,485 Pre-Training PD genes highlighting 32 genes in common, and therefore experiencing editing events. The Genes of Interest in PD were compared to 16,785 Post-Training PD genes identifying 30 genes in common. No significant differences were observed between edited and unedited Genes of Interest in PD genes between Pre- and Post-Training PD samples (p=0.5540). These three gene lists represent genes with known association to PD, have shown differential expression between Pre-and Post-Training PD samples (6), and are being differentially edited between the two groups (S4).

Thirty-two unique Genes of Interest in PD from Pre-Training PD patients and 30 unique Genes of Interest in PD from Post-Training PD patients all showed over-enrichment in Reactome hierarchical pathways including Immune System, Signal Transduction, Gene Expression, Programmed Cell Death, and Circadian Clock. In Pre-Training PD samples, Cell Response to Stimuli pathway was also enriched. Together, these 88 genes identified either in common between Pre- and Post-Training PD samples or as unique to a group (S4) represent a gene list of significance that may play a role in the changes in PD symptomatology, ADAR editing, and editing patterns observed between Pre- and Post-Training groups.

### The number of high or moderate impact editing events occurring in PD genes vary between PD, Control, Pre- and Post-Training PD samples

We further explored whether the total number of ADAR editing events observed in PD genes varied between sample groups. For each subject category, the total editing events included high and moderate impact editing events, along with low impact and modifier editing events. Within Older Male samples, 16,452 total editing events were recorded within PD genes in at least two samples, while there were 18,743 total editing events in PD Males, 16,144 total editing events in Younger Males, 8,408 total editing events in Pre-training PD samples and 6,647 total editing events within Post-training PD samples. Notably, the fractions of high or moderate impact editing events (in other words, those that result in amino acid change with likely functional or structural consequences for affected proteins) also vary across groups. Significant differences were identified through Chi^2^ two-tailed analysis when comparing Older Males to PD Males (p=0.0001), Older Males to Younger Males (p=0.0065), Pre-training PD Males to Post-Training PD Males (p=0.0207), and Older Males to Post-Training PD Males (p=0.0255). Significance was not achieved when comparing the number of high or moderate impact editing events found in PD Males to Younger Male samples (p=0.1829). Together, that the proportion of high and moderate impact editing events in PD Males is more similar to the number of editing events observed in Younger Males than in patients from a similar age group, yet the editing events are occurring in genes associated with different physiological pathways, provides further evidence of dysregulated ADAR editing in PD.

Likewise, when we considered the number of high or moderate impact editing events that resulted in nonsense-mediated decay, significant differences were identified when comparing Older Males to Post-Training PD Males (p=0.0001), with the higher fraction of these editing events in Post-Training category (Fig 2). A similar trend was significant (Chi-square test, p=0.0744). When comparing high or moderate impact editing events occurring in PD genes in protein coding regions, significant differences were identified between Older Males and PD Males (p=0.0482) and Older Males to Post-Training PD Males (p=0.0001). Changes in nonsense-mediated decay regulation have been implicated in a variety of neurodegenerative conditions (82) (83) (84), including PD (85), and thus, the observed differences in editing events may point to candidate genes involved in PD pathogenesis and/or those affected by exercise. Results are shown in Figs 2-4.

**Fig 2:**
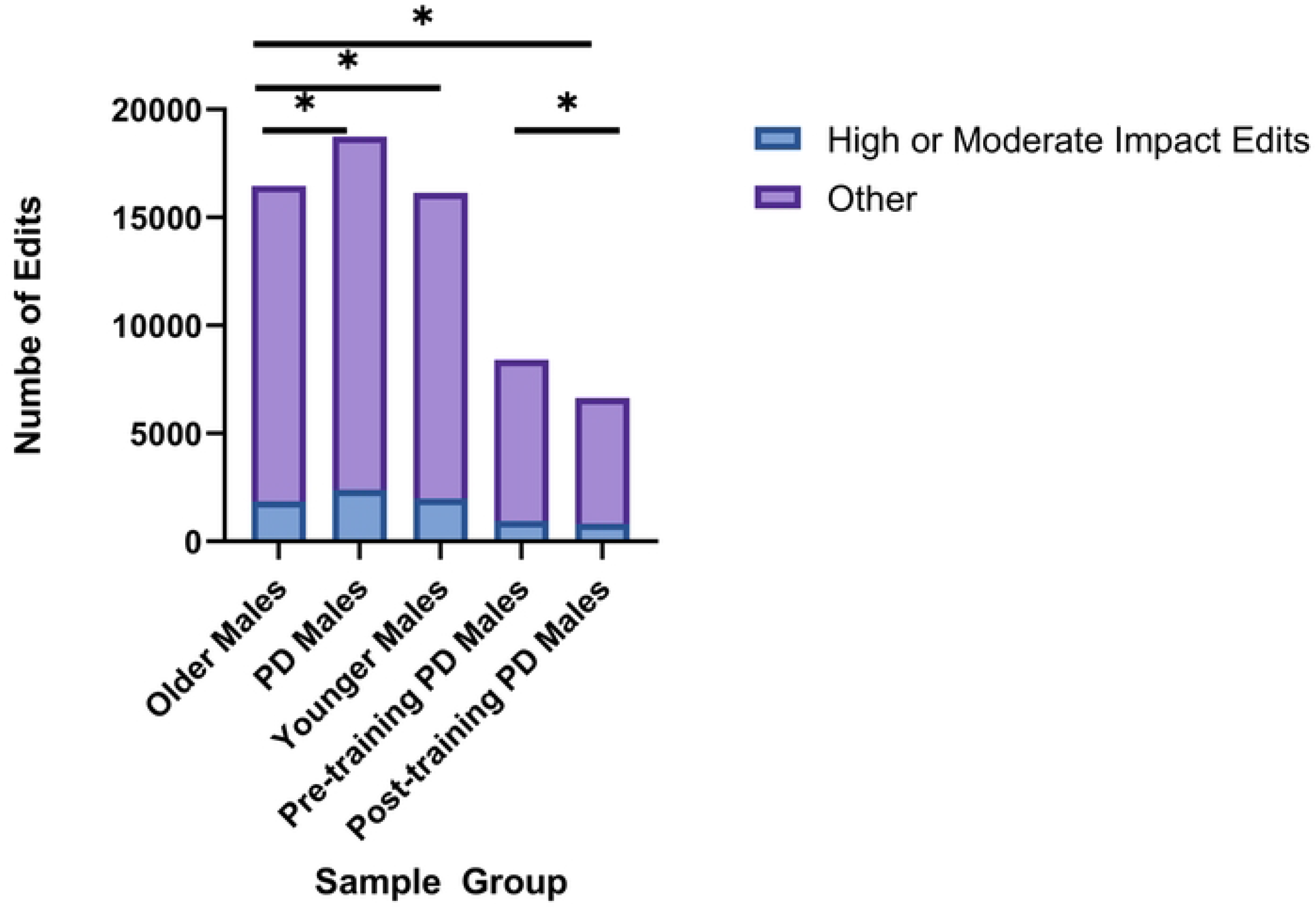
Significant differences in the number of high or moderate editing events between sample groups. Significant differences were observed in the number of high or moderate impact editing events when comparing editing patterns between Older Males and PD Males (Chi-square test, p=0.0001), Older Males and Younger Males (p=0.0065), Pre-Training PD Males and Post-Training PD Males (p=0.0207), and Older Males and Post-Training PD Males (p=0.0255).

**Fig 3:**
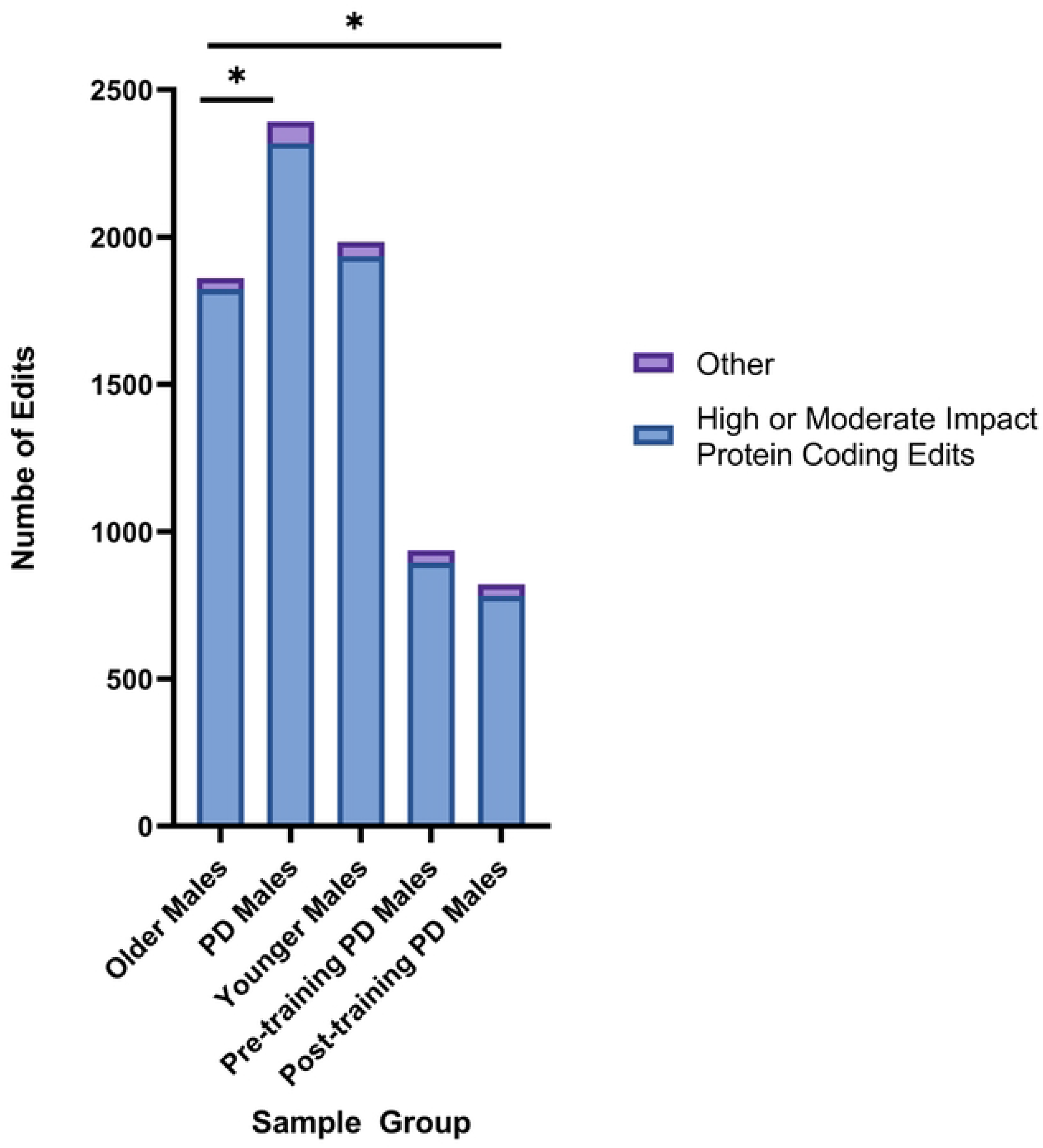
Significant differences in number of high or moderate impact protein coding editing events between sample groups. Significant differences were observed in the number of high or moderate impact protein coding editing events when comparing the number of edits between Older Males and PD Males (Chi-square test, p=0.0482) and Older Males and Post-Training PD Males (p=0.0001).

**Fig 4:**
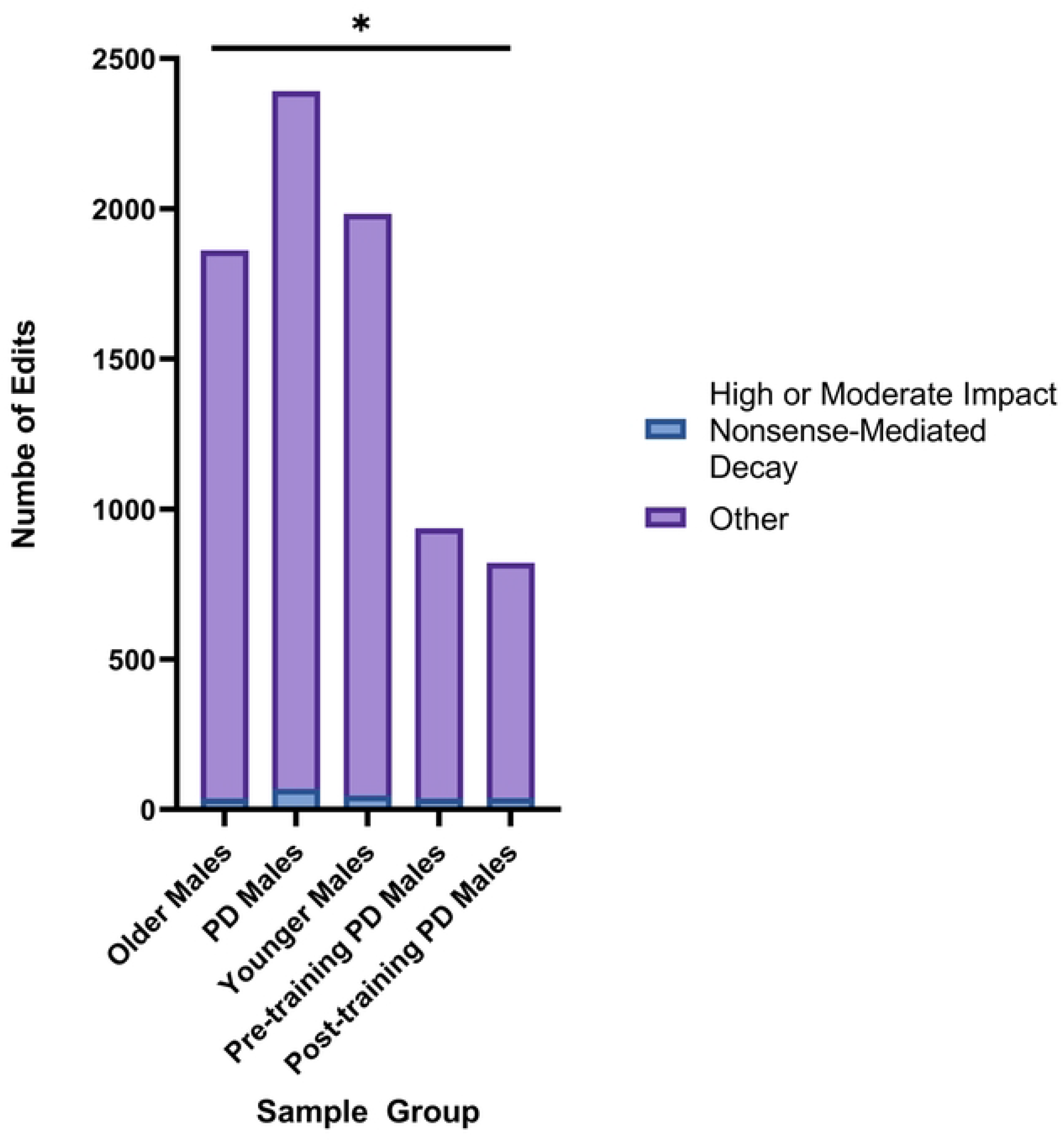
Significant differences in number of high or moderate impact nonsense-mediated decay editing events. Significant differences were observed in the number of high or moderate impact editing events resulting in nonsense mediated decay when comparing the number of edits between Older Males and Post-Training PD Males (Chi-square test, p=0.0001).

### The number of deleterious and tolerated protein-coding editing events occurring in PD genes vary between PD, Control, Pre- and Post-Training PD samples

We further explored the potential impact of ADAR editing on the proteins from the NCBI PD gene list using SIFT (Sorting Intolerant from Tolerant) (77), through SNPNexus annotations (76). We only considered deleterious and tolerated A/G and T/C variants with no dbSNP associations that were observed in two or more samples. The chi-square test was used to analyze the relationship between the amounts of deleterious and tolerated A/G and T/C variants. Significant differences were observed between deleterious and tolerated protein outcomes between Older Males and PD Males (p= 0.0069), Pre-training PD Males and Post-Training PD Males (p=0.0371), and Older Males and Post-Training PD Males (p=0.0001) (Fig 5).

**Fig 5:**
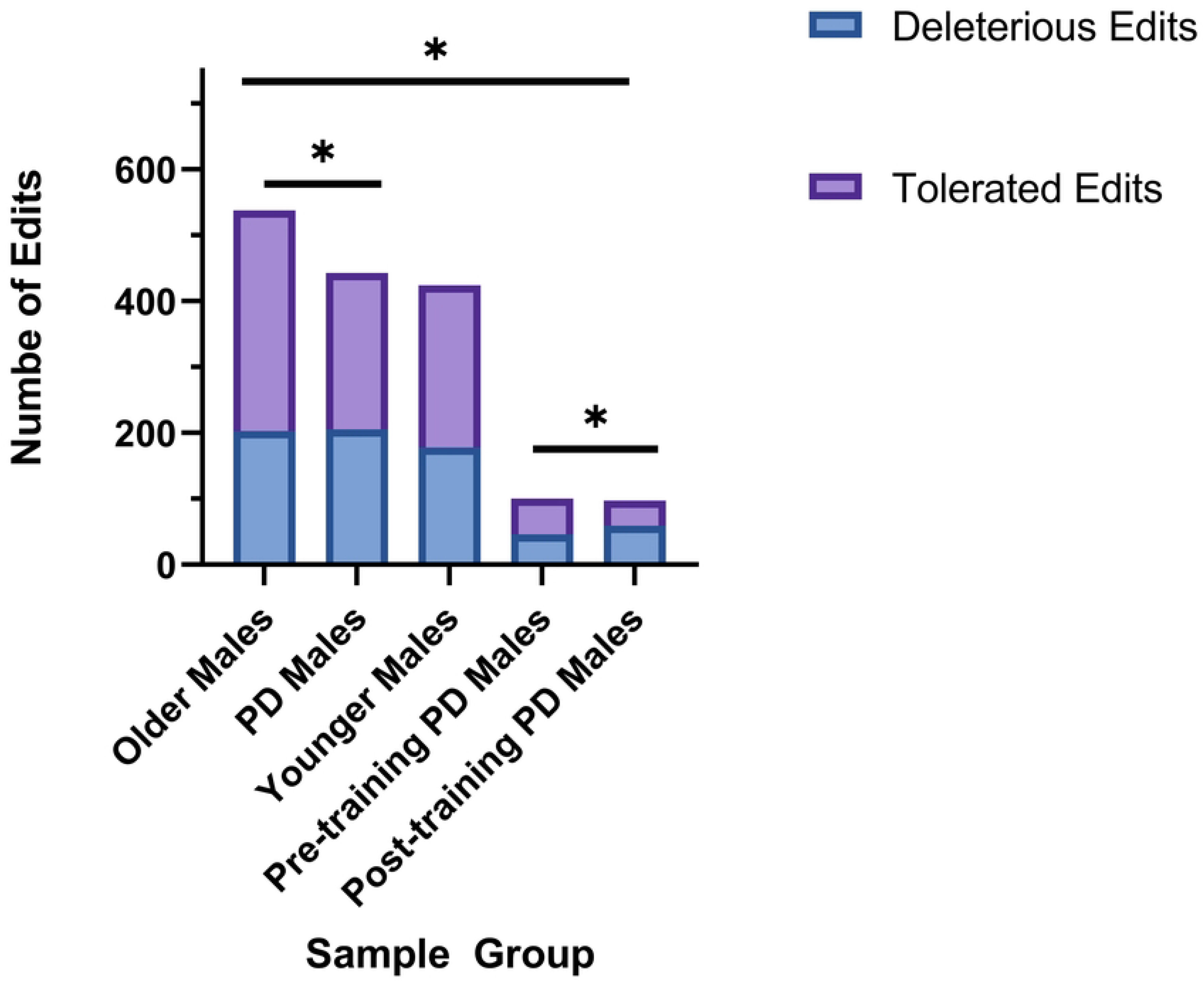
Significant differences in deleterious and tolerated editing events on protein outcomes. Significant differences were observed between deleterious and tolerated protein coding editing events as identified by SIFT when comparing Older Males and PD Males (Chi-square test, p= 0.0069), Pre-training PD Males and Post-Training PD Males (p=0.0371), and Older Males and Post-Training PD Males (p=0.0001).

### Expression of ADAR genes vary between disease state, age, and pre-post training samples

While in the Lavin et al. (6) study ADAR genes were not identified as differentially expressed when the entire transcriptomes were considered, because there is a nuanced non-linear relationship between the levels of ADAR editing and expression of individual ADAR genes (86), we wanted to explore whether any differences in ADAR genes expression can be identified among subject groups. ADAR expression was compared between groups revealing marked differences between groups with significance achieved when comparing PD Males to Post-Training PD Males (t-test, p=0.035). The highest expression of ADAR (ADAR1), which includes the interferon-inducible isoform ADARp150 (87), was seen in PD Male samples with 9.83 average TPM followed by Older Males (average TPM= 9.67) and Younger Males (average TPM =9.45) (Fig 6). The lowest expression of ADAR was demonstrated in Post-training PD Males (average TPM = 7.67). An average of 8.58 TPM was observed for ADAR in Pre-Training PD Male samples. The expression of ADARB1 (ADAR2) was more consistent between Older and PD Males with an average of 3.39 TPM and 3.32 TPM respectively. ADARB1 expression in Younger Males was less than Pre- or Post-Training PD Males with 2.51 TPM in Younger Males, 3.29 TPM in Pre-Training PD Males, and 3.34 TPM in Post-Training PD Males. There was little to no expression of ADARB2 (ADAR2) in any group, as ADARB2 is generally limited to expression in the brain (88) (Fig 6). Global editing events were highest in PD Males (on average, 6,468.2 events per sample) and Post-Training PD Males (6,477.8 events per sample) followed by Pre-Training PD Males (6,360.5 events per sample), Older Males (6,284.1 events per sample) and Younger Males (5,902.6 events per sample) (Fig 7).

**Fig 6:**
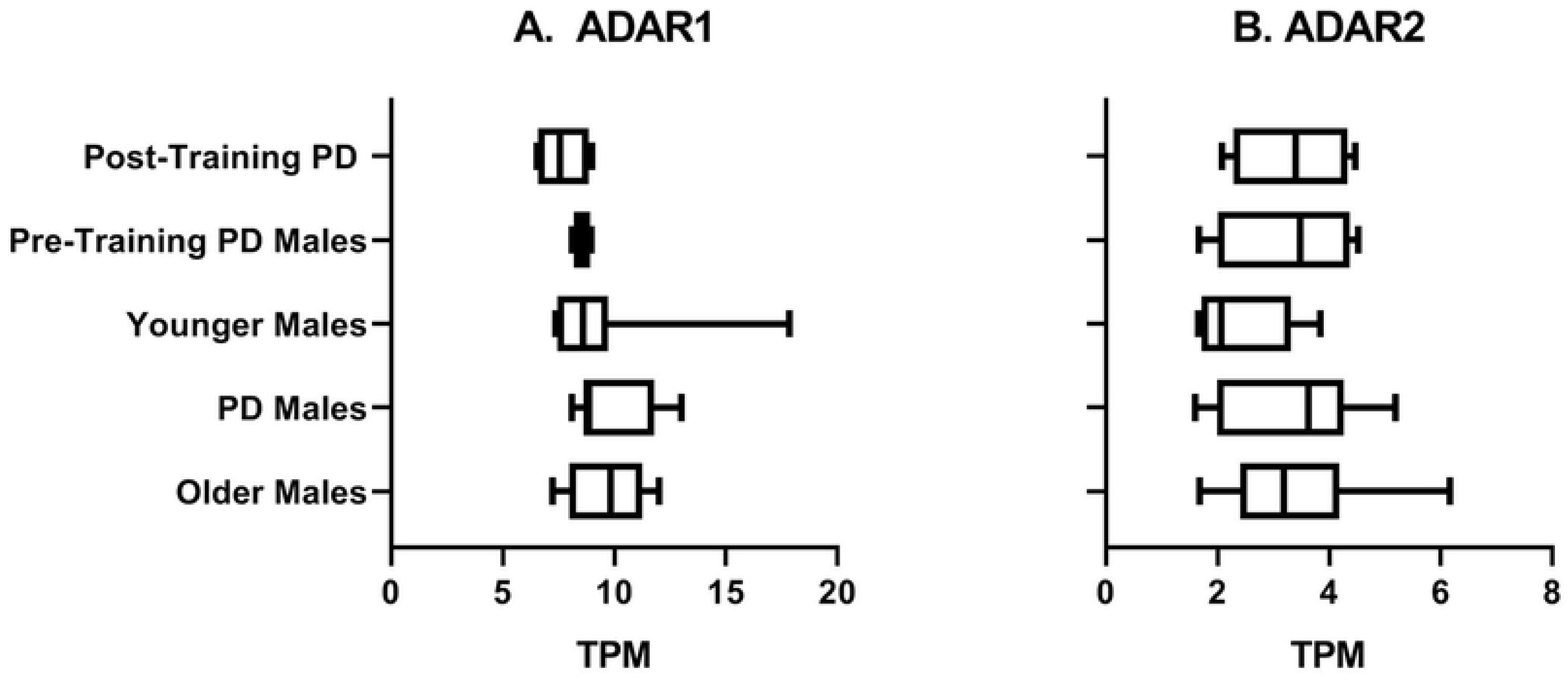
ADAR genes expression levels shown by age, disease state, and training status. (a) ADAR (ADAR1) and (b) ADARB1 (ADAR2) expression, in transcripts per million (TPM), varied somewhat between sample groups, although the pairwise differences did not reach statistical significance (Krushkal-Wallis tests, p values > 0.05). ADARB2 (ADAR3) expression (not shown) was minimal in all sample groups as ADARB2 is primarily expressed in the brain.

**Fig 7:**
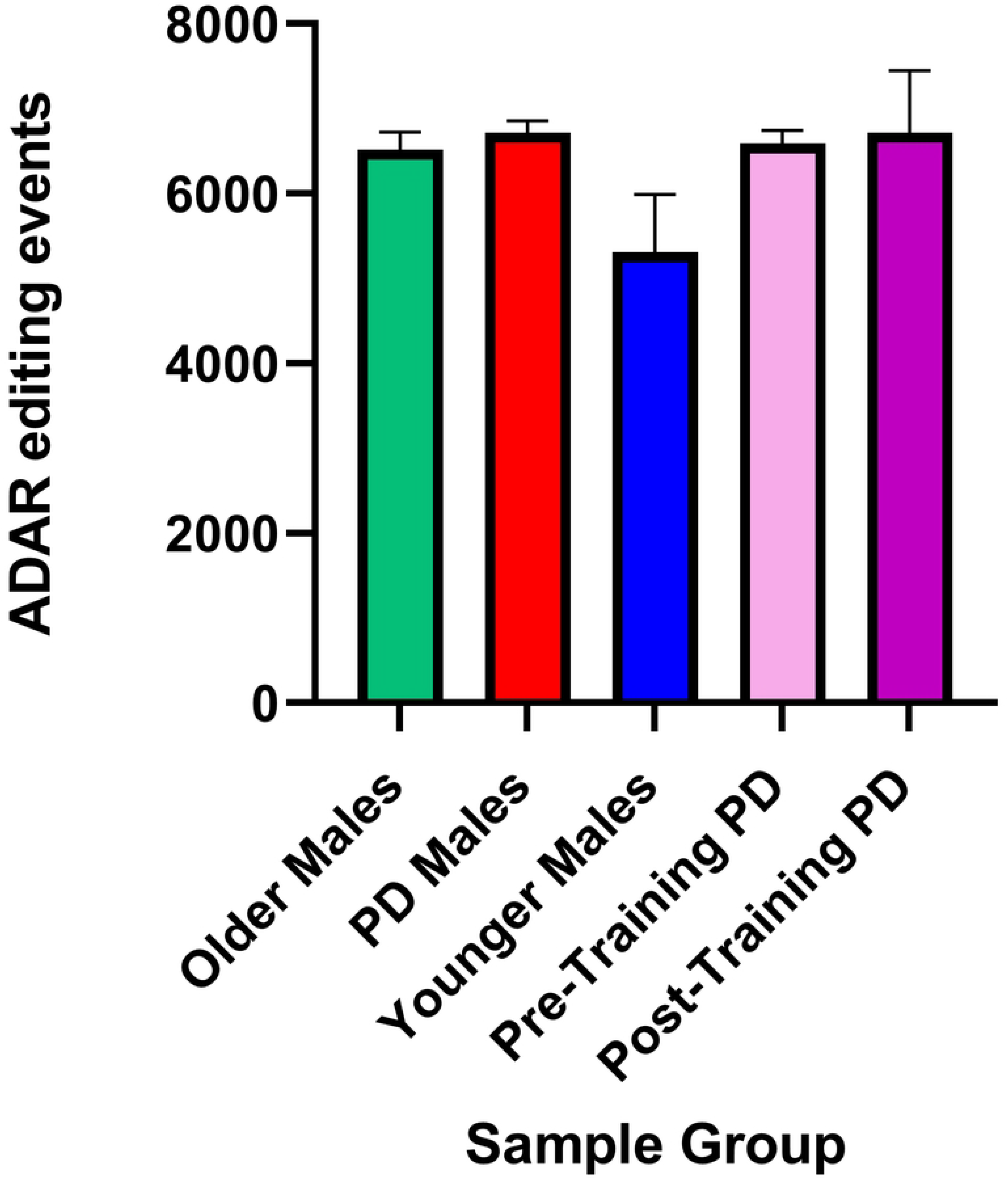
The number of putative ADAR editing events compared between sample groups. Although the mean number of A/G or T/C editing events varied by the group, with the highest number of events identified in PD Males and the lowest number of events observed in Younger Males, none of the pairwise comparisons were statistically significant (Mann-Whitney test, p value > 0.05).

## Discussion

Our results extend Lavin et al. (6) findings of differentially expressed genes to the differences in RNA (ADAR) editing patterns between subject categories, suggesting that exercise training in PD may have effects both on gene expression and differences in RNA/ADAR editing patterns, thus, potentially underlying functionally relevant downstream changes in protein expression and function. In addition, we find that editing patterns change across age groups, and are affected by PD status. From our analysis, we propose the need to further scrutinize the genes in which variations in editing patterns are observed for altered protein function in relation to their editing events and further propose the need for such studies in larger populations and expanded populations that include females.

Specifically, we observed a shift in high and moderate impact editing when comparing samples by age, disease state, and training status. When comparing the pathways associated with enriched PD genes with high or moderate impact edits between Older and Younger Males, editing within Cytokine Signaling in the Immune System and Metabolism of Lipids pathways seem to take precedence. On the other hand, in Younger Males, more PD genes with high or moderate impact edits are associated with Vesicle Mediated Transport. When comparing Older Males to their diseased counterparts, Developmental Biology and Metabolism of Lipids are enriched in Older Males and are not in PD Males. PD genes functioning in the Innate Immune System, Generic Transcription, Gene Expression, RNA Polymerase II Transcription, and Axon Guidance pathways seem to experience more high and moderate impact ADAR editing in Pre-Training PD Males than genes from the post-training samples.

While this study showed that high and moderate impact editing events in PD genes decrease with aging in this analysis, the trend is not observed when examining the total numbers of putative ADAR editing events. The numbers of A/G and T/C editing events are highest in PD and Post-Training PD samples (on average, 6,468.2 and 6,477.8 editing events, respectively) followed by Pre-Training PD Males and Older Males (6,360.5 and 6,284.1 editing events, respectively), with the lowest number identified among Younger Males (5,902.6 editing events), although none of the pairwise comparisons were statistically significant (Mann-Whitney test, p > 0.05) (Table 1). When changes following the exercise were considered, the pattern was inconclusive, with half the patients showing 10+% increase in the number of identified editing events Post-Training, and the other half showed the opposite trend. However, this is based on a very small sample of only four patients.

Likewise, ADAR1 shows elevated expression in PD Males (9.83 TPM) followed by Older Males (9.67 TPM) and Younger Males (9.45 TPM). As inflammation levels are known to increase with age (89,90), it is not surprising that ADAR1 expression is somewhat higher in Older Males than Younger Males; however, we see even greater elevation in PD. The two seemingly contradictory findings of a decrease of high and moderate impact editing with age, yet an increase in ADAR expression suggest that not only changes in editing enzymes are significant between ages and disease, but also that the way that genes are being edited – such as locations and/or level of specific editing events within transcripts - may have downstream significance.

A major limitation of this analysis is the absence of subjects’ health histories, medication profiles, and/or knowledge of any co-morbidities that may be present in addition to PD, although the study participants were screened and excluded in the case of specific diagnoses. For example, many of the participants were taking medications related to PD or other pharmaceutical drugs such as cyclooxygenase (COX) inhibitors/NSAIDs (7) which may affect ADAR expression/editing. This medication history may in turn influence whether the extent of editing went up or down post-exercise (S Fig 2). However, further studies are necessary to substantiate this effect. As the majority of the samples were procured from individuals of advanced age, the likelihood of comorbidities is elevated (91) and may ultimately affect the functional pathways identified as enriched. The ability to adjust for these factors lies outside the range of this study. In addition, only four subjects had both Pre- and Post-training samples available, thus limiting the power of our analysis. Additional uncertainty can be attributed to unknown differences in sample handling and preparation. Some samples had depth of a less than 50 million reads (S2), thus, some editing events may have been missed during variant calling. Moreover, inferences of protein functional outcomes due to nucleotide changes, made by computational tools such as SnpEff and SIFT, may also underestimate the extent of downstream consequences. Lastly, the wide range of PD phenotypes observed in this disease give credence to hypotheses suggesting a broad range of genetic and environmental factors that may contribute to subcategories of PD manifestation (59,60); however, the number of samples prevents us from considering PD subcategories.

Importantly, albeit consistent with the bulk of PD research to-date, the lack of female samples creates an obstacle in furthering our understanding of PD, its progression, and how sex impacts its manifestation. The Lavin et al. study (6) included only 10 female samples: 3 Older Females, 3 PD Females, 3 Younger Females, and 1 Post-Training PD Female. This sample number was not robust enough to make valid comparisons between groups and our previous findings that ADAR editing may vary between the sexes (62), so we are not including these analyses along with this research. However, we did conduct a basic analysis of female data, with some intriguing results.

Significant differences in the number of high or moderate impact editing events in PD genes were observed when comparing Older Males to Older Females (p=0.0001), PD Males to PD Females (p=0.0001), Younger Males to Younger Females (p=0.0001), and Post-Training Males to one Post-Training Female (p=0.0001) (Fig 8). These pronounced differences suggest that relationships between PD pathology and ADAR editing may be drastically different between the sexes. Many have noted the phenotypic differences in PD between males and females (92–94). A recent study by Sandor et al. (60) noted gender differences in PD, specifically when considering a specific PD phenotype denoted as ‘Axis 2’ suggesting not only gender influences, but also an interplay between sex and multifactorial genetics. We propose the urgent need to further scrutinize RNA editing patterns in PD, the specific genes involved in significant pathways identified in this study, and the variations in how transcripts are edited between the sexes and among the range of PD phenotypes.

**Fig 8:**
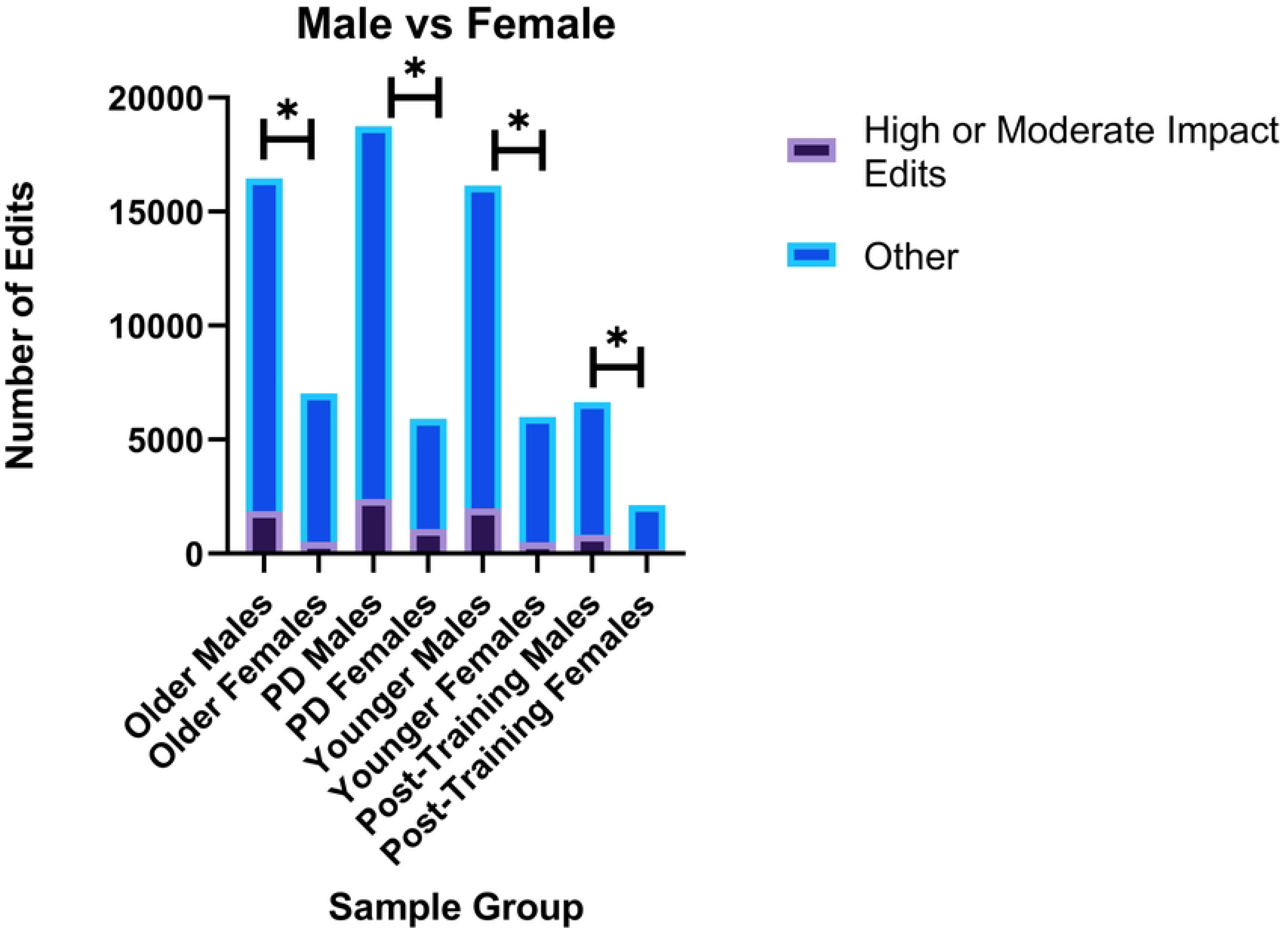
Comparison between the number of high or moderate impact editing events between males and females. Significant differences were observed in the number of high or moderate impact editing events when comparing Older Males vs Older Females (Chi-square test, p=0.0001), PD Males vs. PD Females (p=0.0001), Younger Males vs. Younger Females (p=0.0001), and Post-Training PD Males vs one Post-Training PD Female (p=0.0001).

## Conclusion

Here, we have shown that alterations in ADAR editing patterns between PD, Control, and Post-Training PD male samples may lead to changes in protein functions, which then may contribute to the manifestation and progression of PD pathology. Our findings also show some intriguing changes in ADAR editing patterns that occur post-exercise, although further studies are needed to delineate the role of RNA editing in therapeutic exercise. Likewise, further functional studies are needed for a set of PD-related genes that we identified as harboring high or moderate impact editing events and/or inferred deleterious protein outcomes. Critically, we emphasize the need to further elucidate the role of ADAR editing in PD progression in females, as impact of sex is largely omitted in PD studies to-date.

Ultimately, while mutations in specific candidate genes have been suggested as culprits in PD pathogenesis, no monogenic targets have been definitively identified, making it more likely that a variety of genetic and environmental influences are at play in the manifestation of the neuroinflammation indicative of PD. ADAR editing, a known factor in the pathology of multiple neurodegenerative and psychiatric disorders (28,64,80), must be acknowledged as a potential role-player in PD pathology. Further research must strive to understand how dynamic changes in ADAR editing may function as a causal agent in PD progression or whether RNA editing dysregulation is simply an outcome of the neuroinflammation and neurodegeneration present in the disease.

## Acknowledgements

This study was partially supported by the LaunchPad Award from Healthy Communities Research Institute (Kent State University).

---

**S Fig 1:**
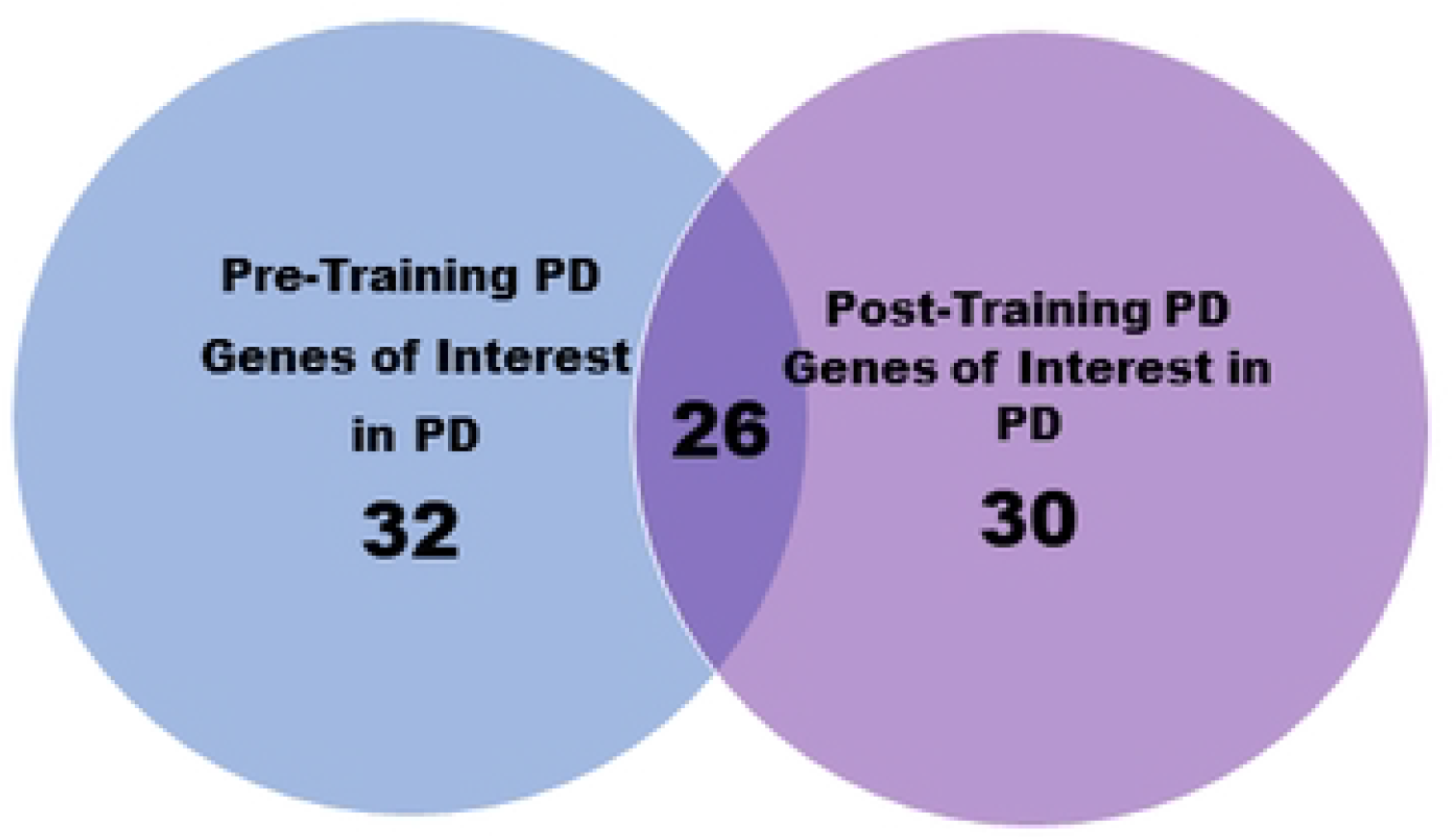
Pre- and Post-Training PD genes found in common with the Genes of Interest in PD List and between groups. Unique PD genes experiencing ADAR editing events were identified within Pre-Training PD and Post-Training PD samples and compared with the Genes of Interest in PD Gene List. The Genes of Interest in PD List includes genes in common between the NCBI PD Gene List and the genes shown to be differentially expressed between pre- and post-training PD patients in skeletal muscle (6).

**S Fig 2:**
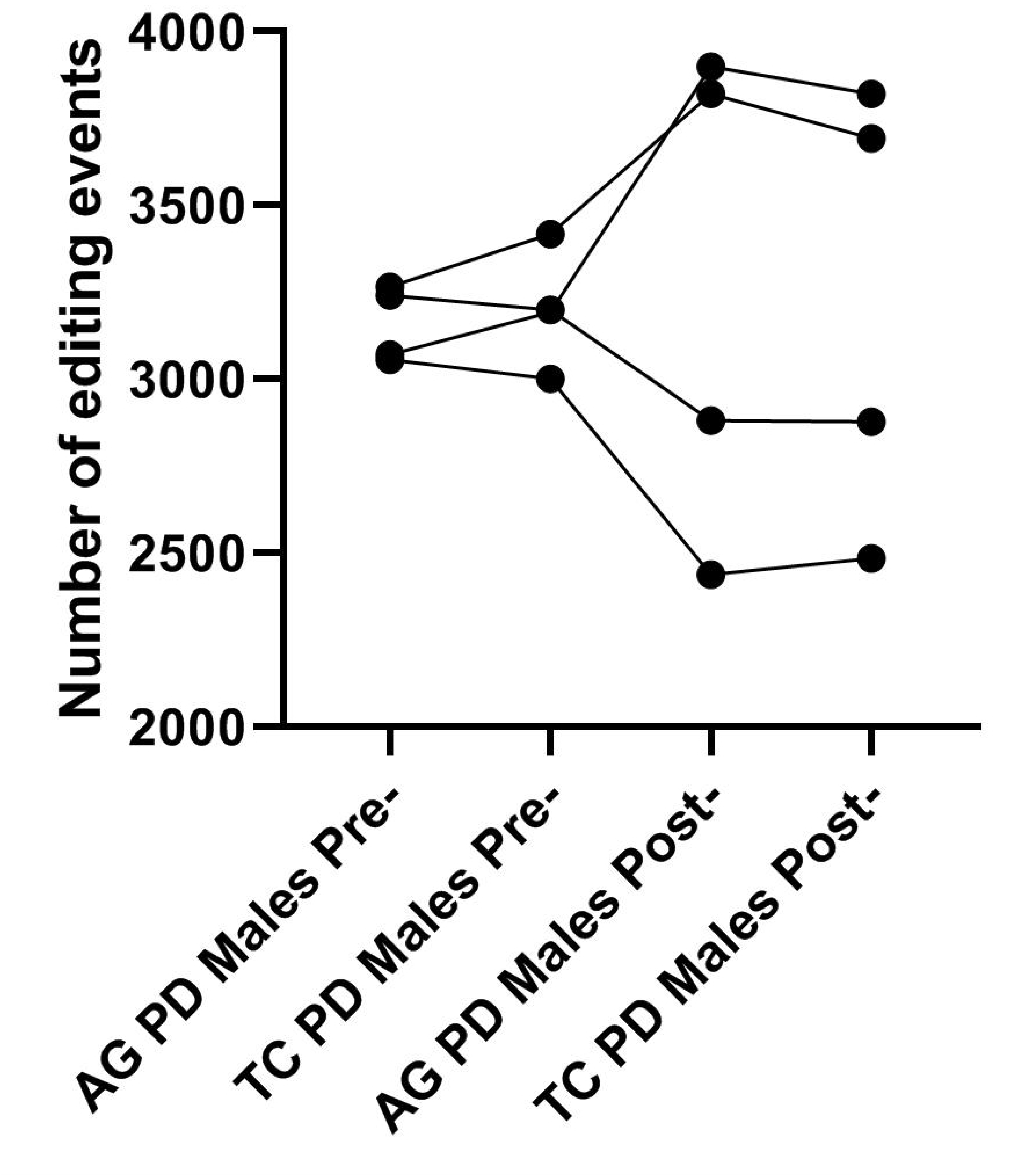
Changes in the number of A-to-G and T-to-C editing events PD genes from matched Pre- and Post-Training samples. Each dot represents counts of editing events, with lines connecting dots from individual PD patient pre- and post-exercise.

